# Subtype-Specific Dependencies and Drug Vulnerabilities Enable Precision Therapeutics in Head and Neck Cancer

**DOI:** 10.64898/2025.12.05.692684

**Authors:** Joel Vaz, Songli Zhu, Mateo Useche, Lauren Shih, Emily Marchiano, Slobodan Beronja, Bruce E Clurman, Cristina Rodriguez, Brittany Barber, Marina Chan, Taranjit S Gujral

## Abstract

Molecular heterogeneity in head and neck squamous cell carcinoma (HNSCC) is well recognized, yet existing subtype frameworks remain largely descriptive and have not translated into therapeutic decision-making. Here, we establish a mechanistic platform that converts transcriptomic diversity into drug-actionable tumor states. Integrating multi-cohort RNA-seq from 727 tumors across five independent datasets, genome-scale CRISPR dependency maps, and pharmacologic screening, we define distinct tumor survival circuits across HPV-negative HNSCC and nominate subtype-matched therapeutic strategies. These circuits encompass a proliferative axis (MYC, MET/FAK, inflammatory and translational programs), an epithelial-differentiated/adhesion program, an EMT-like state with stromal activation, and mitochondrial/oxidative metabolic states, each mapping to selective liabilities (e.g., mitotic/autophagy control, ERBB/PI3K and cadherin signaling, OXPHOS/mitochondrial translation, and G2/M-integrin-Notch pathways, respectively). We then develop a transcriptomic predictor of EGFR-inhibitor response using machine learning and validate it in prospectively collected, fresh patient-derived 3D microtumors. The resulting 13-gene signature identifies erlotinib-responsive tumors (R = 0.93) and maps biologically to an epithelial-differentiated state, outperforming EGFR expression alone. Our study establishes a subtype-to-dependency-to-therapy framework, enabling precision stratification and providing a clinically feasible path for prospective biomarker deployment.

## INTRODUCTION

Head and neck squamous cell carcinoma (HNSCC) remains a major global health burden, with ∼600,000 new cases annually and limited progress in survival, particularly among human papillomavirus (HPV)-negative tumors, where five-year survival still hovers near 50% despite multimodal therapy^1^. Over the past decade, large-scale genomic and transcriptomic efforts have revealed substantial biological heterogeneity in HNSCC, identifying discrete molecular programs and refining tumor classification beyond anatomic site^2–7^. These studies have defined canonical subtypes, including classical, basal, mesenchymal, and atypical/HPV-related groups, and provided foundational insights into disease biology. However, these subtype frameworks have largely remained descriptive. They illuminate tumor diversity but do not yet provide a mechanistic basis for therapy selection, nor do they accurately predict clinical response or guide patient stratification.

The critical challenge is therefore not simply to catalogue heterogeneity, but to operationalize it. Despite extensive molecular profiling, HNSCC has lagged behind other solid tumors in realizing precision oncology. Actionable genomic drivers are uncommon; common alterations such as TP53, CDKN2A loss, and NOTCH pathway mutations lack targeted therapies; and current biomarkers, including PD-L1 expression and tumor mutational burden, incompletely predict response to immunotherapy^8,9^. Likewise, although EGFR is highly expressed in HNSCC and EGFR inhibitors have been evaluated clinically, therapeutic benefit remains modest and inconsistent, underscoring the inadequacy of single-marker approaches and the need for biological context.

Here, we explore how HNSCC molecular subtypes relate to functional dependencies and therapy response. We integrate five independent RNA-sequencing cohorts to generate one of the largest unified transcriptomic atlas of HNSCC to date and derive reproducible molecular states across HPV-positive and HPV-negative disease. We then superimpose genome-scale CRISPR gene-essentiality maps and unbiased pharmacologic screening data to uncover survival circuits and subtype-specific therapeutic liabilities. This integrative strategy resolves the well-known basal and classical categories into mechanistically distinct tumor states, distinguished by proliferative signaling, epithelial differentiation, immune-stromal composition, and metabolic wiring, transforming prior descriptive labels into functional tumor identities with clear biological underpinnings and therapeutic implications.

Finally, we operationalize these biological states to create a clinically implementable prediction framework. Leveraging subtype-informed transcriptomic features, we build and rigorously cross-validate a machine-learning model for EGFR-inhibitor response, then prospectively test and validate this model in fresh, patient-derived 3D microtumors. This direct demonstration that molecular predictors trained in silico can forecast therapeutic sensitivity in primary human tumor tissue represents a key conceptual shift, moving beyond descriptive associations toward a practical precision-oncology framework in which tumor state informs biological dependency and, ultimately, treatment response.

Together, this work provides a principled path to translate tumor heterogeneity into therapeutic decisions in HNSCC. More broadly, it offers a generalizable blueprint for converting multi-omic cancer taxonomies into functional states that guide rational drug selection, model-informed clinical trial design, and prospective biomarker deployment across solid tumors where molecular subtypes are known but not yet actionable.

## RESULTS

### Defining molecular tumor sates in HNSCC through multi-cohort transcriptomic integration

To comprehensively characterize the molecular landscape of HNSCC, we integrated RNA-seq data from five independent cohorts, FHCC, GSE74927, GSE205308, GSE178537, and TCGA^10^, yielding a total of 753 samples (**Fig. 1A, Tables S1 and S2**). Raw count data from TCGA were downloaded via the GDC portal, while all other raw files were obtained from the NCBI GEO database. After alignment using STAR^11^, samples with fewer than 60% uniquely mapped reads were excluded, resulting in a final dataset of 727 samples. To account for differences across data sources, we aligned and quantified gene expression (**Fig. S1A**), then applied batch correction using ComBat-Seq^12^. UMAP visualization after correction showed well-mixed samples from different cohorts, indicating successful harmonization (**Fig. 1B**). Next, we identified the optimal number of clusters by comparing Silhouette and Davies-Bouldin scores, both of which supported *k* = 9 (**Fig. 1C**). The Silhouette score measures how well each sample fits within its assigned cluster versus others, with higher values indicating better-defined clusters, while the Davies-Bouldin index evaluates intra-cluster similarity and inter-cluster separation, where lower values suggest more distinct clustering. Unsupervised k-means clustering at *k* = 9 partitioned the dataset into nine distinct subgroups (**Fig. 1D**). Although we also tested alternative methods including HDBSCAN, Gaussian mixture modeling, and hierarchical clustering (**Fig. S1B-S1D**), we selected k-means for its ability to define multiple clusters and its strong support from both evaluation metrics.

**Fig. 1:**
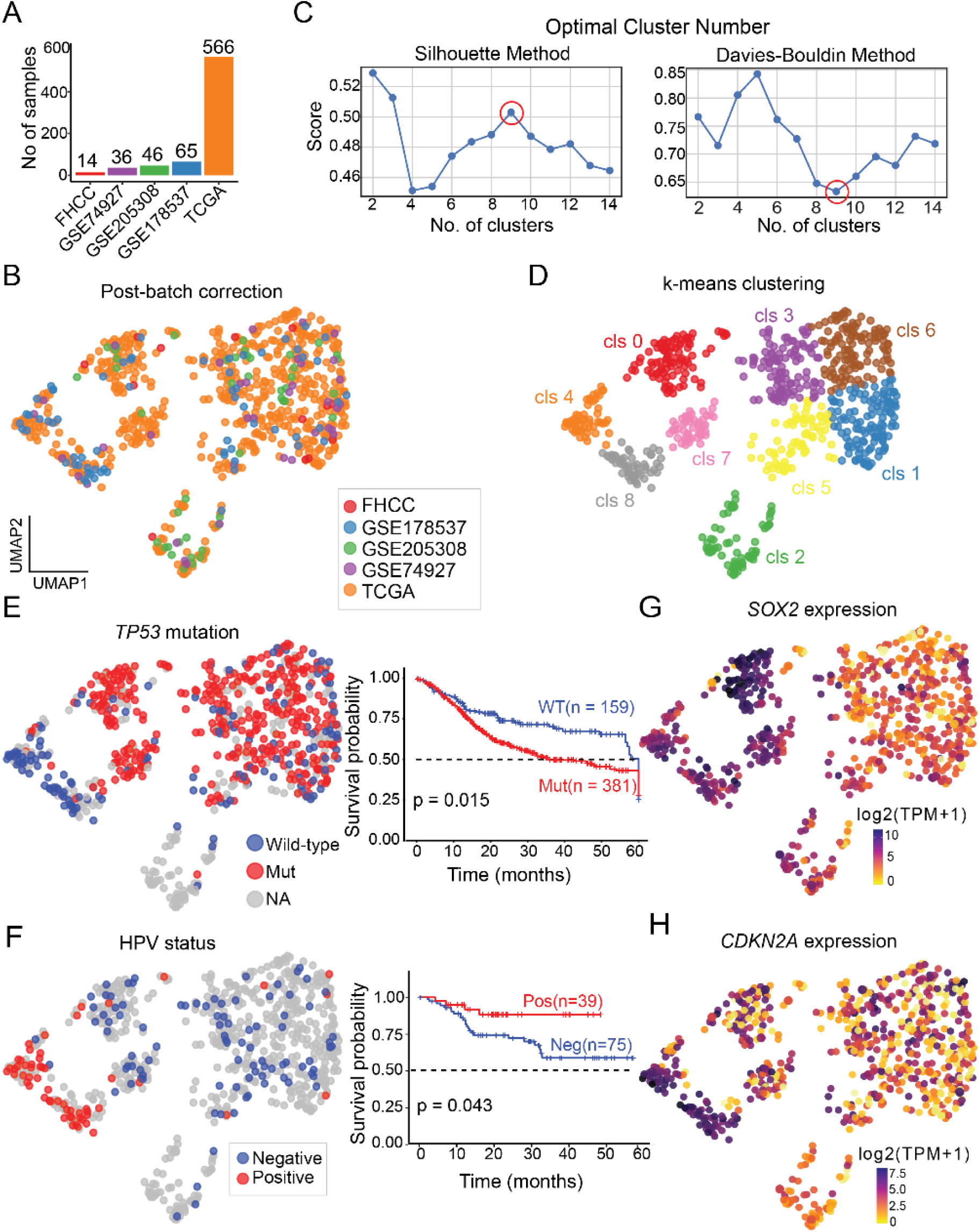
Transcriptomic clustering of HNSCC identifies nine molecular subgroups with distinct genetic and clinical features. **A** Bar plot showing the number of RNA-seq samples collected from different datasets, including Fred Hutch Cancer Center (FHCC), GEO datasets, and TCGA. **B** UMAP visualizing HNSCC samples after batch correction, with colors indicating the dataset of origin. **C** Optimal cluster number determination using the Silhouette (left) and Davies-Bouldin (right) methods. The Silhouette index is maximized, and the Davies-Bouldin index is minimized at k = 9. **D** K-means clustering of the integrated dataset at optimal cluster number, with each color representing a distinct cluster. **E** UMAP showing TP53 mutation status (wild-type vs. mutant), alongside Kaplan-Meier survival curves comparing overall survival between the two groups. **F** UMAP indicating HPV status (negative vs. positive), with the corresponding Kaplan-Meier survival curves for each group. **G** UMAP showing SOX2 log2-TPM expression levels across the samples. **H** UMAP displaying CDKN2A log2-TPM expression levels across the samples.

To explore clinical and molecular features associated with each cluster, we overlaid key annotations onto the UMAP. *TP53* mutation status, available for ∼82% of samples, was enriched across most clusters, except for clusters 4 and 8 (**Fig. 1E** and **Fig. S1E**). As previously reported^13^, patients with *TP53* mutations had significantly poorer survival. HPV status, determined by p16 testing in ∼17% of samples, revealed that HPV-positive tumors localized to distinct regions in the UMAP, particularly in clusters 4 and 8 where *TP53* mutations were less common (**Fig. 1F** and **Fig. S1F**). These tumors, as expected^14,15^, were associated with better survival outcomes. This inverse relationship between HPV positivity and *TP53* mutation was further supported by ISH data (**Fig. S1I**). Smoking history was more broadly distributed but appeared less common in clusters 4 and 8 (**Fig. S1G, S1H**). The Sex-determining region Y-box 2 (SOX2) gene, a key regulator of embryonic stem cell fate, has been associated with improved prognosis in patients with HNSCC^16^. When SOX2 expression was overlaid onto the UMAP, higher expression was observed in clusters 0, 7, 3, and 8 (**Fig. 1G**). In contrast, loss or mutation of *CDKN2A*, a common alteration in HPV-negative tumors^17^ was generally associated with lower expression across most clusters, except for clusters 4 and 8 (**Fig. 1H**), which are enriched for HPV-positive tumors. The mutual exclusivity between HPV positivity and TP53 mutation and/or CDKN2A loss observed in our distinct cluster has been previously described^18^. Cluster 4 predominantly contained normal samples; thus we designated it as the non-tumor cluster. Overall, these findings align with established models of HNSCC pathogenesis, where HPV-positive tumors form distinct molecular subgroups. In our analysis, we identified two clusters among HPV-positive tumors and six distinct clusters among HPV-negative HNSCC.

To investigate the biological underpinnings of the HNSCC subgroups identified through unbiased clustering methods, we performed a multi-step transcriptomic analysis (**Fig. 2A**). First, we conducted differential gene expression (DEG) analysis to identify genes that were significantly up- or downregulated in each tumor cluster compared to all other tumors. Using these DEGs, we generated pre-ranked gene lists for each cluster and applied GSEA-PreRanked^19^ to assess pathway enrichment across multiple gene set collections, including Hallmark, KEGG, Reactome, Biocarta, ImmuneSigDB, and Gene Ontology. Next, we used single-sample Gene Set Enrichment Analysis (ssGSEA) to compute pathway activity scores for each individual tumor sample. These scores were projected into UMAP space to visualize how specific biological programs are distributed across the different tumor subtypes (**Fig. 2A**).

**Fig. 2:**
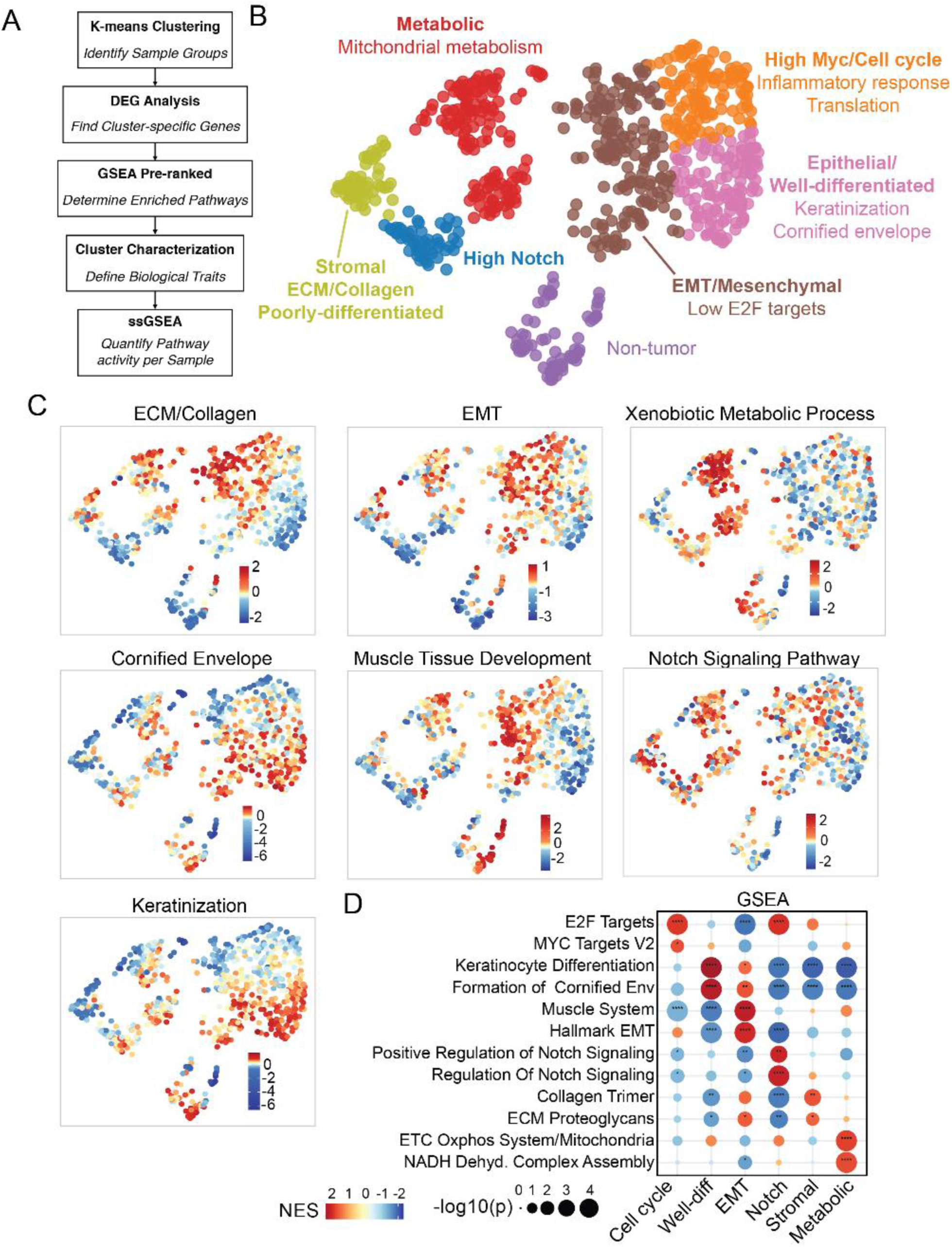
Transcriptomic subtypes of HNSCC reveal distinct biological programs and pathway enrichments. **A** Overview of the workflow. K-means clustering was applied on the integrated RNA-seq dataset to define molecular subgroups. Differential expression analysis was then used to identify cluster-specific genes. Pre-ranked GSEA was applied to determine enriched biological pathways and subtype traits. **B** UMAP visualization of HNSCC samples, color-coded by molecular subtype. Each cluster is annotated with defining biological programs, including metabolic, EMT/mesenchymal, epithelial/well-differentiated, stromal/ECM, and high Notch signaling features. **C** UMAP plots showing single-sample GSEA enrichment scores (z-score normalized) for representative pathways, including ECM/collagen remodeling, EMT, xenobiotic metabolism, cornified envelope, muscle tissue development, Notch signaling, and keratinization. **D** Bubble plot summarizing average pathway enrichment across subtypes (from GSEA Pre-ranked). Bubble size reflects significance (–log10 p-value) and color corresponds to normalized enrichment score (NES). Significance levels are derived from permutation-based test. ns: p ≥ 0.05, *: p < 0.05, **: p < 0.01, ***: p < 0.001, ****: p ≤ 0.0001

Among the two HPV-positive clusters, both exhibited active Notch signaling; however, they were distinguished by differences in stromal composition. Cluster 0 showed strong Notch pathway activity, enrichment in cornified envelope genes, and low stromal signatures, features consistent with a predominantly epithelial phenotype (**Fig. 2B**). In contrast, Cluster 8 also displayed Notch signaling but was enriched in stromal-related pathways, including extracellular matrix (ECM) remodeling (**Fig. 2B**), indicating a more mesenchymal-like or immune-infiltrated microenvironment.

Among the six HPV-negative clusters, we identified distinct biological subtypes based on pathway enrichment patterns. Cluster 6 was marked by strong cell cycle and MYC signaling, along with activation of MET/FAK signaling, inflammatory pathways, and components of the translational machinery; features indicative of a highly proliferative and potentially immune-modulated subtype (**Fig. 2C**). In contrast, Cluster 1 exhibited signatures of a well-differentiated epithelial phenotype, including enrichment in keratinocyte differentiation, epidermal development, cornified envelope formation, and translation-related pathways (**Fig. 2C**). Together, Clusters 1 and 6 were grouped into a ‘highly proliferative, well-differentiated module’. Cluster 0 and 7 show strong enrichment in mitochondrial metabolism and oxidative phosphorylation pathways, suggesting a possible dependence on energy production; we refer to this as ‘MT metabolism-enriched module’. While both clusters are linked to metabolic activity, only Cluster 0 showed enrichment of Wnt signaling. (**Fig. 2C**). Clusters 3 and 5 showed enrichment in mesenchymal and muscle-related processes, such as epithelial-to-mesenchymal transition (EMT), muscle development, and myogenesis. These clusters also exhibited low expression of E2F target genes, consistent with reduced cell cycle activity and increased potential for migration or invasion, leading us to classify them as the ‘EMT module’. Thus, based on pathway enrichment and underlying biological processes, the six HPV-negative clusters were grouped into four distinct modules: a highly proliferative, well-differentiated module; an MT metabolism-enriched module; an EMT module; and an immune-modulated proliferative module.

### Relationship to prior subtyping frameworks

Our refined HPV-negative tumor states build on and extend the TCGA subtypes^7,20^. Consistent with prior work, we recover basal, mesenchymal, and classical patterns. However, our integrated analysis separates the TCGA basal group into two mechanistically distinct states, a proliferative MYC/MET-FAK-high program and a well-differentiated epithelial program and subdivides the classical class into two mitochondrial-metabolism states. The mesenchymal class aligns directly with our EMT-enriched state. These refinements provide greater biological resolution than traditional labels and clarify distinct therapeutic vulnerabilities embedded within the basal and classical groups.

### Tumor microenvironment signatures distinguish molecular subtypes

Having identified four distinct modules among HPV-negative tumors and two among HPV-positive tumors, we next sought to characterize the contribution of specific cell types, particularly stromal and immune cells, across these subtypes. We performed cellular deconvolution using xCell^21^, a gene signature-based method that estimates the relative enrichment of 64 immune and stromal cell types in bulk tissue transcriptomes (**Table S3**). This analysis revealed varying degrees of infiltration by epithelial cells, fibroblasts, macrophages, T cells, and other immune populations across the molecular subgroups (**Fig. 3A**). As expected, the HPV-positive subtypes showed increased enrichment of immune cells, including CD4+ and CD8+ T cells, consistent with an “immune subtype” classification^22^. Among these, the stromal-enriched HPV-positive cluster showed higher fibroblast scores and lower enrichment of epithelial cells and keratinocytes, suggesting a more mesenchymal-like tumor microenvironment.

**Fig. 3:**
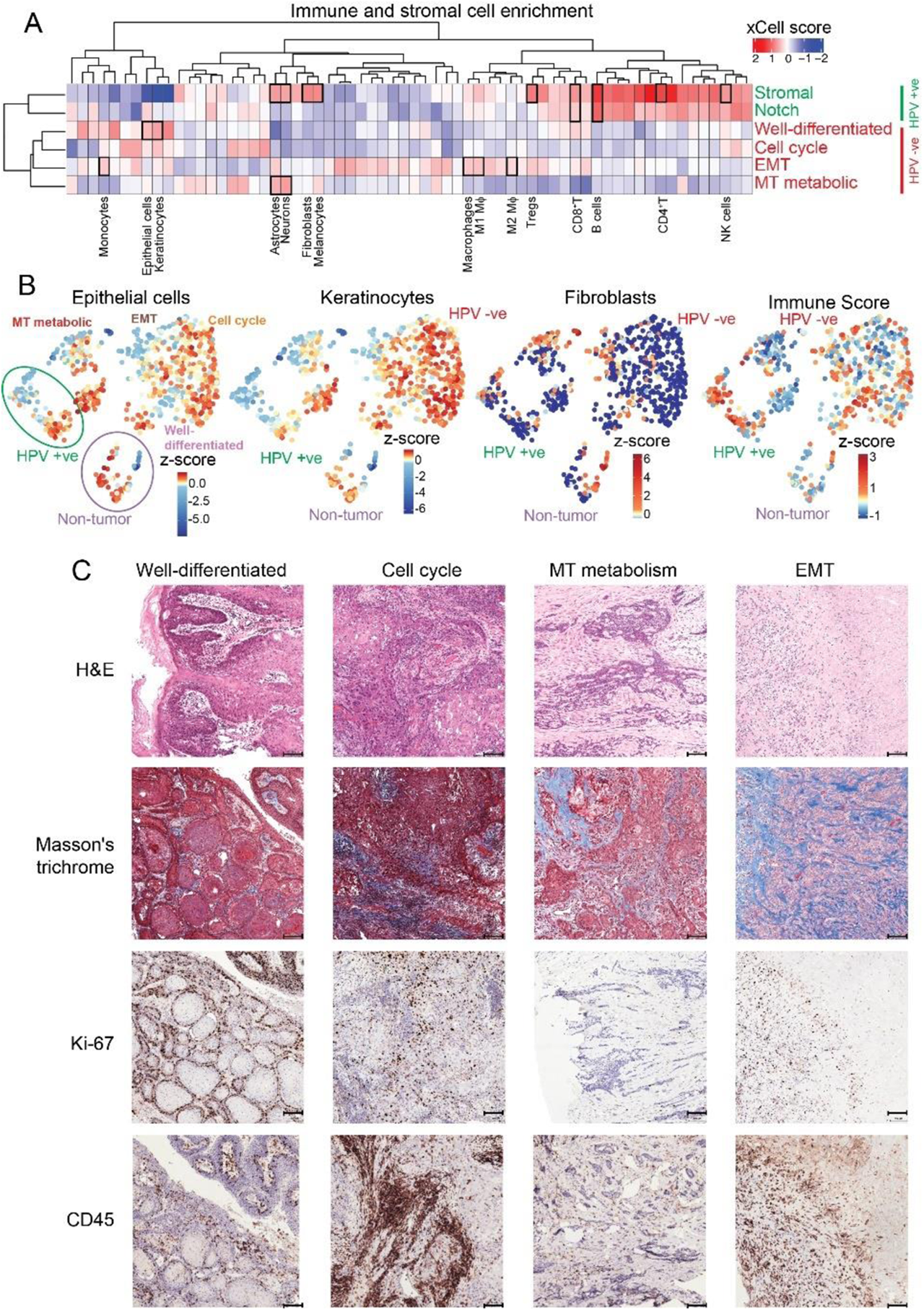
Immune and stromal landscapes distinguish molecular subtypes of HPV-Negative HNSCC. **A** Heatmap of z-normalized xCell scores for 64 immune and stromal cell types across molecular subgroups of HNSCC. Rows correspond to subtypes (MT metabolic, EMT, cell cycle, well-differentiated, Notch, stromal, HPV+), and columns correspond to inferred cell populations. Red indicates higher enrichment, blue indicates lower enrichment. **B** UMAPs illustrating the predicted enrichment of selected cell types and immune scores at the single-sample level. Each point is colored by the inferred abundance of epithelial cells, keratinocytes, fibroblasts, CD8+ T cells, or overall immune score. **C** Representative histology and immunostaining of HPV-negative SCC tissues from each major subtype. Top: H&E staining shows overall tissue morphology. Second row: Masson’s Trichrome staining highlights subtype-specific collagen deposition. Third row: Ki-67 staining indicates proliferative activity. Bottom: CD45 staining shows patterns of immune cell infiltration, with differences across well-differentiated, cell cycle, mitochondrial metabolism, and EMT subtypes.

Among the HPV-negative tumors, the well-differentiated and cell cycle-driven modules displayed higher enrichment of epithelial cells and keratinocytes, reflecting a more epithelial-like phenotype (**Fig. 3B**). In contrast, the EMT module showed elevated levels of M2-like macrophages, the highest among all subtypes, indicating a strongly immunosuppressive and pro-invasive microenvironment. Interestingly, the mitochondrial metabolism-enriched module exhibited low enrichment of most immune cells, epithelial cells, and keratinocytes, but showed relatively higher enrichment of astrocyte- and neuron-like cell types, suggesting a unique stromal composition in this subgroup.

Next, we sought to evaluate cell type composition and stromal characteristics using histological analysis of freshly collected HPV-negative HNSCC samples. We performed H&E, Masson’s Trichrome, Ki67, and CD45 staining to assess tissue architecture, ECM content, proliferation, and immune cell infiltration. (**Fig. 3C**). The well-differentiated subtype showed organized epithelial structures and uniform nuclear morphology, consistent with an epithelial-rich phenotype. The cell cycle-enriched subtype exhibited densely packed nuclei, reflecting high proliferative activity. In contrast, the mitochondrial metabolism and EMT-enriched subtypes displayed spindled cell morphology and reduced cell-cell adhesion, features associated with poor differentiation and a mesenchymal-like state. Masson’s Trichrome staining revealed subtype-specific ECM patterns. Collagen (blue) was sparse and stromally localized in well-differentiated tumors. In the cell cycle-enriched subtype, collagen was variably distributed within cellular regions, suggesting intermediate stromal remodeling. EMT-enriched tumors showed dense, disorganized collagen bundles throughout, indicative of a fibrotic microenvironment. Ki67 staining confirmed high proliferation in the cell cycle-enriched subtype, while mitochondrial metabolism-enriched tumors showed sparse Ki67 positivity. CD45 staining indicated abundant immune infiltration in the cell cycle-enriched subtype and minimal infiltration in the mitochondrial metabolism-enriched subtype, highlighting contrasting immune landscapes. Overall, these findings confirm that the observed differences in cell type composition and stromal features align with the predicted subtypes, supporting the validity of our classification through independent histological analysis.

### Subtype-specific genetic dependencies in HNSCC

To explore potential therapeutic vulnerabilities associated with HPV-negative HNSCC subtypes, we aimed to identify representative *in vitro* models with drug response data (**Fig. 4A**). The Cancer Cell Line Encyclopedia (CCLE) provides a comprehensive resource of molecularly characterized cancer cell lines, including drug sensitivity profiles, making it a valuable tool for identifying subtype-specific dependencies^23^. We began by calculating single-sample GSEA (ssGSEA) scores for 26 available HNSCC cell lines in CCLE using the top 50 upregulated and downregulated pathways previously identified for each cluster. We then computed pairwise cosine similarities between these cell line ssGSEA profiles and those from our integrated tumor dataset (**Fig. 4B, Table S4**). The resulting similarity matrix revealed that certain cell lines closely resembled subtypes characterized by cell cycle activity e.g. *SCC25, HSC3, PECAPJ49*, whereas others aligned with metabolic e.g. *SNU1041, DETROIT562, FADU, SNU1214, A253*, or EMT-associated pathways e.g. *BICR56, BICR22, PECAPJ15, BICR6, HSC2*. Based on these profiles, we selected a representative panel of 18 HPV-negative cell lines, ensuring that each subtype was included (**Fig. 4B**).

**Fig. 4:**
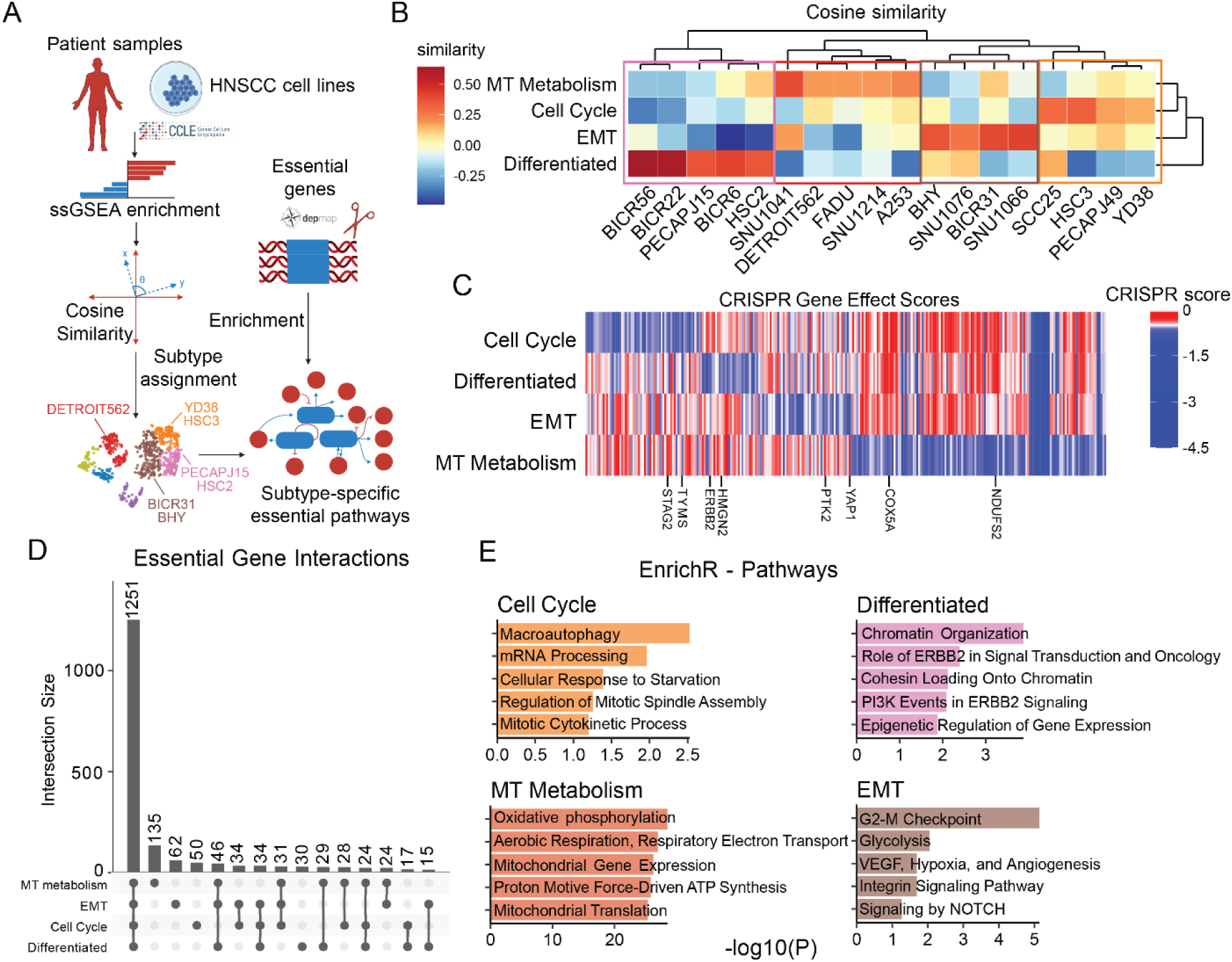
CRISPR Dependency mapping reveals subtype-specific vulnerabilities in HPV-negative HNSCC. **A** Schematic overview of the CRISPR-based analysis used to map cell lines to the pathways essential for their survival. **B** Heatmap of cosine similarity scores between tumor subtype profiles and HNSCC cell lines. Red indicates higher similarity, blue indicates lower similarity, with clustering highlighting correspondence between subtypes and representative lines. **C** Heatmap of CRISPR gene effect scores across subtypes. Red indicates non-essential genes, blue indicates strong essentiality, with subtype-specific dependencies such as STAG2, TYMS, ERBB2, and YAP1 highlighted. **D** UpSet plot of essential gene intersections across HPV-negative subtypes. The largest shared set (n=1251) reflects core survival genes, while smaller subsets represent subtype-specific vulnerabilities. **E** EnrichR pathway enrichment analysis of subtype-specific essential genes. Distinct pathways were associated with each subtype.

To identify subtype-specific dependencies, we utilized CRISPR-based gene effect scores from DepMap, which quantify the degree to which a gene is essential for cell survival. These scores are derived from CRISPR-Cas9 loss-of-function screens, where individual genes are knocked out and the impact on cell viability is measured. A lower gene effect score indicates higher dependency, meaning the gene is critical for the survival of that cell line. A cutoff of z-score < -0.5 is typically used to define essential or genes important for cell growth, distinguishing them from non-essential ones (**Fig. 4C**). We leveraged this data to identify essential genes for each HNSCC subtype. We defined an essential gene for a given subtype as one that was essential across nearly all representative cell lines, corresponding to at least 75-80% of members (all or all but one). Intersection analysis showed that while some essential genes were shared across subtypes, others were uniquely associated with specific subtypes. For example, *STAG2* and *TYMS* were found to be important in the cell cycle-enriched subtype, *ERBB2* and *HMGN2* in the well-differentiated subtype, *YAP1* and *PTK2* in the EMT subtype, and *COX5A* and *NDUFS2* in the mitochondrial metabolism-enriched subtype.

Next, we examined the overlap of essential genes across HPV-negative HNSCC subtypes using an UpSet plot (**Fig. 4D**). A large, shared core dominated the landscape (n = 1,251; 84.2% of all essentials), indicating that, beyond subtype-specific features, these tumors converge on a common viability core dependency. Enrichment of this pan-subtype set highlighted fundamental gene-expression and cell-division machinery, including RNA metabolism and processing (mRNA splicing and transport; rRNA processing), ribosome biogenesis and translation (formation and function of 40S/60S subunits; initiation/elongation/termination), DNA replication and repair, and cell-cycle control (G1/S and G2/M transitions, checkpoints, spindle assembly, cytokinesis) (**Table S5**). Together, these pathways define a housekeeping program centered on transcription, RNA processing, ribosome function, protein synthesis, and orderly cell-cycle progression. While a subset was specific to head and neck disease (n = 106), most genes were essential across all cell lines (n = 1,145; **Fig. S2A**). We also observed subtype-restricted essentials: mitochondrial-metabolism (n = 133), EMT (n = 62), cell-cycle (n = 50), and differentiated (n = 30). Although the total number of CRISPR-defined essentials per cell line was broadly similar across groups, the differentiated subtype showed markedly fewer essentials shared among its lines, suggesting greater functional heterogeneity within this class. These data underscore the coexistence of a common survival core with separable, subtype-specific liabilities that motivate targeted follow-up analyses.

To further investigate these subtype-specific dependencies, we performed pathway enrichment analysis using EnrichR on the uniquely essential genes of each subtype (**Fig. 4F**). In the cell-cycle subtype, enriched terms included macroautophagy, mRNA processing, cellular responses to starvation, and regulation of mitotic spindle assembly/cytokinesis. Together, these point to a proliferative state that buffers biosynthetic demand and energy stress (autophagy/starvation programs) while remaining dependent on core mitotic machinery. The differentiated subtype was enriched for ERBB2-mediated PI3K signaling, chromatin organization, and cohesin loading, consistent with an epithelial program that maintains lineage identity and cohesion and relies on growth-factor signaling. The mitochondrial-metabolism subtype showed strong enrichment for oxidative phosphorylation, aerobic respiration, mitochondrial gene expression/translation, and proton motive force-driven ATP synthesis, highlighting a mitochondria-centered energy dependence as a key vulnerability. Finally, the EMT subtype was enriched for the G2-M checkpoint, glycolysis, VEGF/hypoxia/angiogenesis, integrin signaling, and NOTCH signaling, indicating a mesenchymal, microenvironment-interactive state that retains checkpoint control, reroutes metabolism, and leverages stromal/angiogenic and adhesion pathways for viability and invasion. Together, these analyses reveal distinct biological processes underlying each subtype and offer potential therapeutic entry points tailored to their specific vulnerabilities.

### Pharmacologic vulnerabilities across molecular tumor states

To investigate pharmacologic sensitivities linked to HPV-negative HNSCC subtypes, we leveraged the PRISM Repurposing drug screen^24^ across representative cell lines and observed subtype-specific response patterns that mirrored the dominant biological pathways of each group (**Fig. 5A**).

**Fig. 5:**
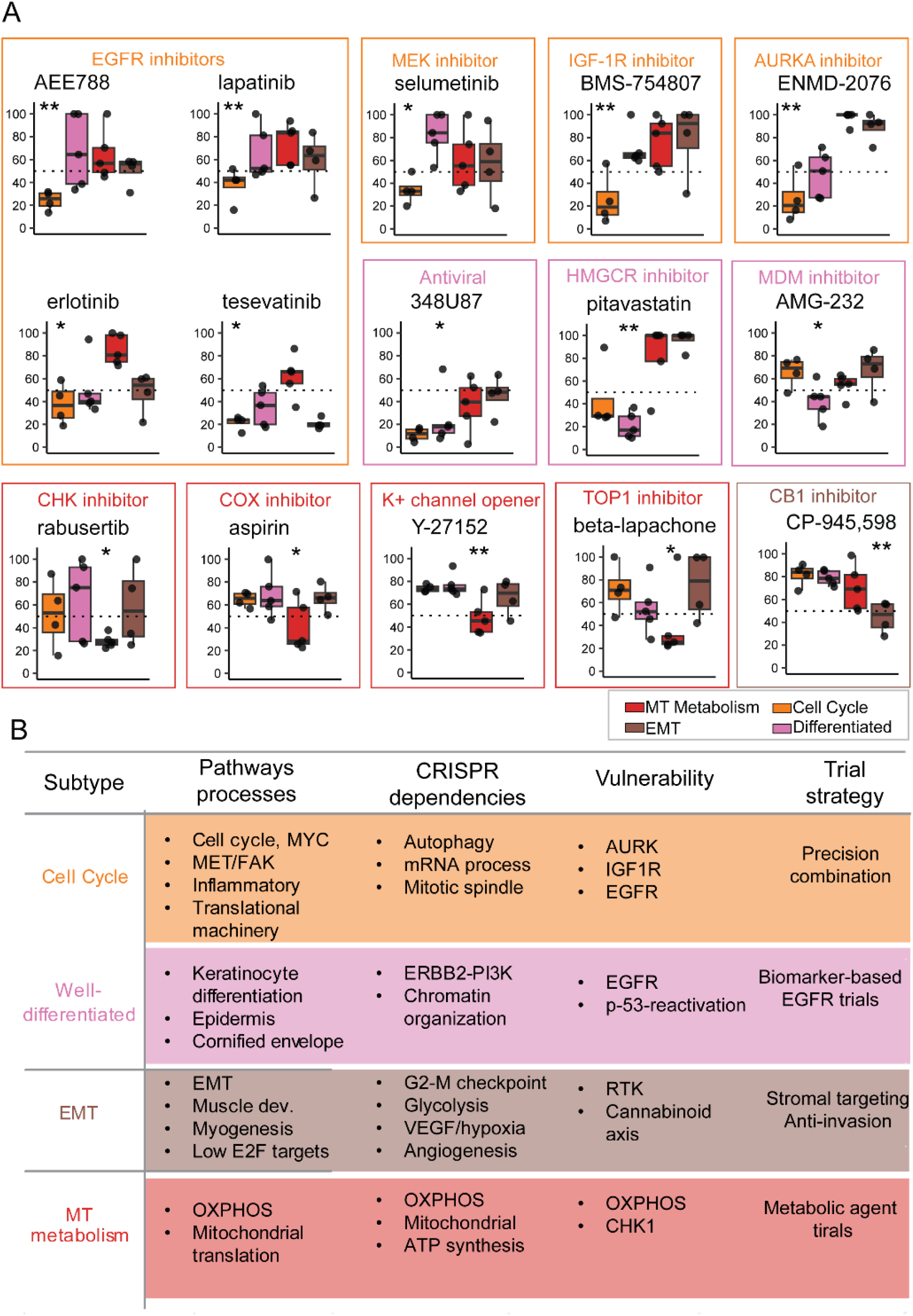
Subtype-specific drug sensitivities reveal distinct therapeutic vulnerabilities in HPV-Negative HNSCC. **A** Boxplots show cell viability after treatment with representative compounds, grouped by subtype assignment. **B** Summary table linking molecular subtypes of HPV-negative HNSCC to their defining biological programs, enriched pathways, candidate therapeutic agents and potential trial strategy.

As an example, EGFR inhibitors, including AEE788, lapatinib, and erlotinib exerted potent cytotoxicity in both the Cell Cycle and Differentiated subtype cell lines. This dual efficacy likely reflects and a dependency on upstream growth factor signaling in highly proliferative cell cycle lines (**Fig. 4E**) and EGFR/ERBB pathway activation in differentiated lines^25,26^. Tesevatinib was the only EGFR inhibitor with notable activity in the EMT subtype.

Consistent with their mitotic dependence (**Fig. 4E**), cell cycle subtype lines also showed pronounced sensitivity to the MEK inhibitor selumetinib, the IGF-1R inhibitor BMS-754807^27^, and the Aurora A kinase inhibitor ENMD-2076^28^, all of which reduced median cell viability to below 40%. These results highlight the vulnerability of this group to disruption of both mitotic and upstream proliferative signaling. In contrast, several agents exhibited selective efficacy in the well-differentiated subtype cell lines. The antiviral compound 348U87^29^, the HMG-CoA reductase inhibitor pitavastatin, and the MDM2 inhibitor AMG-232^30^ were particularly effective in these cells, reducing median viability to 15-45%. These sensitivities align with enrichment of p53 signaling and apoptotic pathways in this subtype, suggesting that disruption of sterol metabolism or p53 reactivation may be promising therapeutic strategies.

The mitochondrial metabolism-enriched subtype cell lines displayed unique sensitivity to compounds that target metabolic stress and redox pathways. Specifically, the CHK1 inhibitor rabusertib, COX inhibitor aspirin, potassium channel opener Y-27152^31^, and the TOP1 inhibitor β-lapachone^32^ each reduced viability below 50% only in this group. These findings are consistent with the subtype’s strong enrichment for oxidative phosphorylation and mitochondrial translation pathways^33^.

Finally, the EMT subtype demonstrated selective vulnerability to the CB1 receptor antagonist CP-945,598^34^, which reduced median viability to ∼45%. This suggests a potential and previously unappreciated role for endocannabinoid signaling in supporting the survival of mesenchymal-like tumor cells. Together, these results demonstrate that drug response profiles recapitulate and expand upon the genetic dependencies identified in earlier CRISPR screens. These findings offer a robust preclinical framework for matching HNSCC molecular subtypes to rational, targeted therapeutic strategies (**Fig. 5B**).

### Predicting EGFR inhibitor response using transcriptomic signatures and patient-derived microtumors

Building on the potential drug links in HPV-negative HNSCC subtypes, we next focused on EGFR inhibition, since EGFR is a commonly overexpressed receptor tyrosine kinase in HNSCC and a clinically actionable target (e.g., cetuximab)^35^. EGFR drives downstream MAPK/PI3K pathways that regulate proliferation and survival^36^. However, clinical benefit with EGFR inhibitors is inconsistent, implying that transcriptional context, rather than EGFR levels alone, shapes sensitivity (**Fig. 6A**). While erlotinib (an EGFR inhibitor) showed stronger responses in HNSCC cell lines compared to all other cancer types (p=<0.001), its effects on viability still varied widely within HNSCC, providing a clear rationale to build a model that predicts response from gene-expression features (**Fig. 6B**)

**Fig. 6:**
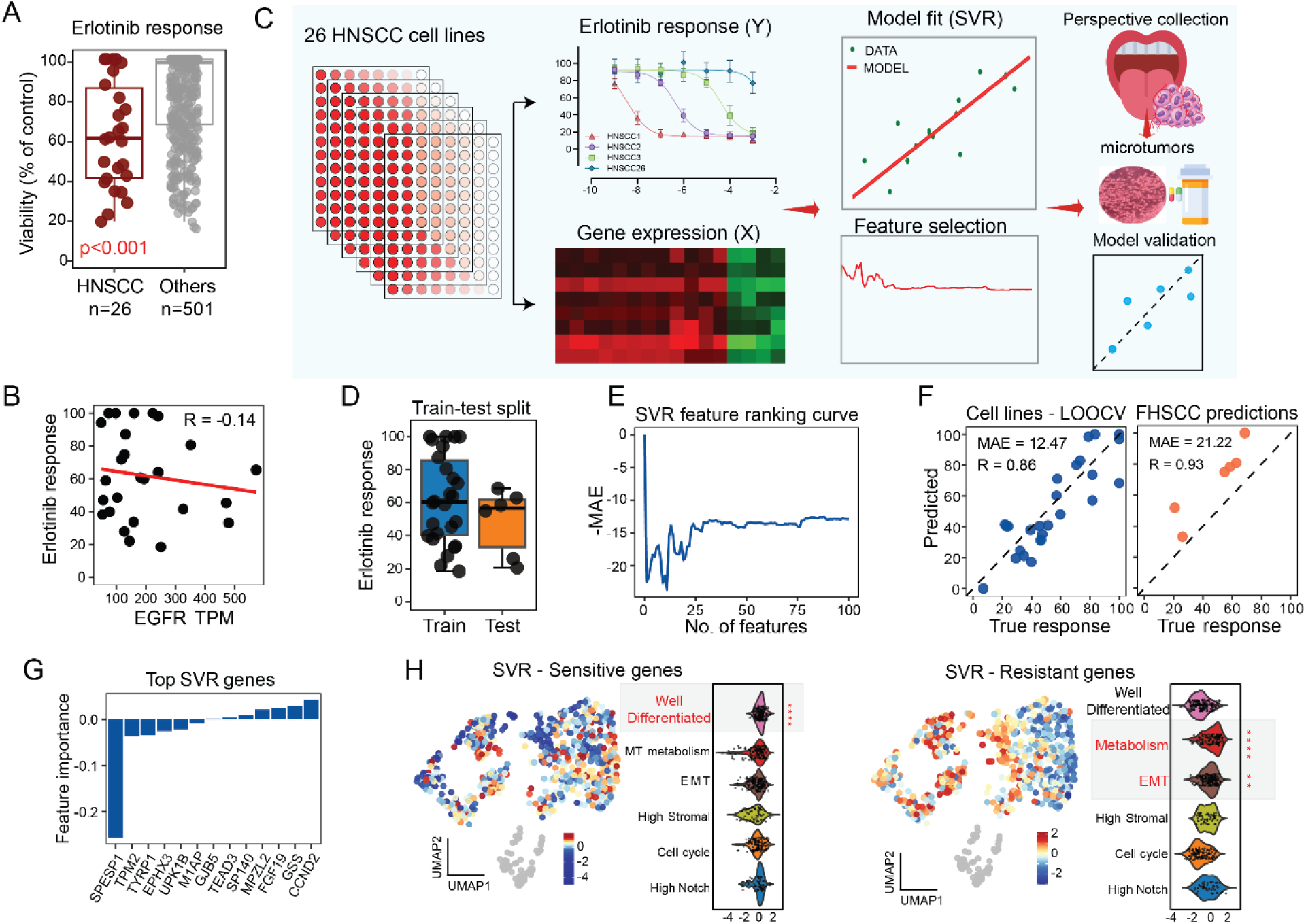
Machine learning identifies gene signatures predictive of EGFR Inhibitor response. **A** Box plot of erlotinib response showing reduced viability in HNSCC lines compared to other cancer cell lines. **B** Scatter plot of erlotinib response versus EGFR mRNA expression (TPM), showing no strong correlation (R = –0.14). **C** Schematic of the supervised learning workflow. RNA-seq from 26 HNSCC CCLE lines was used to train a support vector regression (SVR) model of erlotinib response, which was then applied to predict responses in patient-derived tumors. **D** Box plots of erlotinib response in training (cell lines) and test (patient-derived microtumors) sets, showing reduced viability in clinical samples. **E** SVR feature ranking curve showing cross-validation mean absolute error (MAE) relative to the number of features selected. **F** Scatter plots comparing observed versus predicted responses for cell lines (left, LOOCV; MAE = 12.47, R = 0.88) and patient-derived microtumors (right; MAE = 18.44, R = 0.88). **G** SVR feature importance coefficients highlighting the top predictive genes driving the erlotinib response model. **H** UMAP visualization and violin plots of ssGSEA z-scores for sensitivity-associated (positive coefficient) genes, showing subtype-specific enrichment. Right, UMAP visualization and violin plots of ssGSEA z-scores for resistance-associated (negative coefficient) genes, with significant enrichment across specific subtypes.

To identify patients who are more likely to benefit from anti-EGFR therapy, we developed an approach that integrated erlotinib response data from HPV-negative cell lines (training set, n=26) with their baseline transcriptomics profiles (**Fig. 6C**). We applied a supervised learning framework using support vector regression (SVR) to learn the mapping between gene-expression features and drug response. The best-performing model was then applied to new clinical samples from FHCC cohort (test set, n=6) to evaluate its accuracy and ability to predict anti-EGFR inhibitor response using transcriptomics data alone (**Fig. 6D**).

To identify features predictive of erlotinib response, we employed a recursive feature elimination with leave-one-out cross-validation (RFE-LOOCV) framework using SVR, as previously described in Kang et al., 2022, ^37^. After removing genes with low expression and variance, we subsampled 90% of the training data and performed RFE to rank informative genes. The top 100 ranked genes were evaluated across all cell lines using LOOCV, and the optimal model was achieved using 13 genes, resulting in a minimum cross-validation MAE of 12.47 (**Fig. 6E**).

The true test of the model is its ability to predict drug response in clinical samples. To this end, we recruited 6 HPV-negative HNSCC cases from the FHCC cohort and evaluated the response to erlotinib using patient-derived 3D microtumors. Microtumors offer important advantages over 2D cell lines because they retain the native tumor microenvironments, preserve multiple cell types and can be prepared immediately from surgical specimens^38^. We have recently shown that drug responses in 3D microtumors outperforms sensitivity testing in 2D cultures^38^. For each case, microtumors were generated and treated with erlotinib at 2.5uM and response was measured using real-time glo, as described previously^39^. In parallel, gene expression profiles from these 6 cases were used to evaluate the *in silico* prediction accuracy of our model. Our model demonstrated strong generalization to the test set, achieving an MAE of 21.22 and a Pearson correlation (R) of 0.93 in 6 independent clinical samples (**Fig. 6F**). The final 13-gene predictive signature comprised both sensitivity-associated (*GJB5, TEAD3, SP140, FGF19, GSS, CCND2*) and resistance-associated (*SPESP1, TPM2, TYRP1, EPHX3, UPK1B, M1AP*) features (**Fig. 6G**). Application of this signature to the integrated tumor transcriptomic landscape, using single-sample enrichment scoring, revealed that the sensitivity-associated gene set was significantly enriched in the well-differentiated subtype (Wilcoxon one-sided rank-sum test, *p* < 0.0001), indicating that elevated expression of these genes may underlie heightened erlotinib responsiveness (**Fig. 6H**). In contrast, the resistance-associated signature showed preferential enrichment in the mitochondrial metabolism-enriched subtype, suggesting that upregulation of these genes may contribute to the intrinsic erlotinib resistance observed in this subgroup.

Several of the selected genes are biologically relevant to cell state regulation. *CCND2*, identified in our top predictive gene set, has been observed to be upregulated in less aggressive, more differentiated squamous epithelial tissues and in contexts of *CCND1* downregulation, suggesting a compensatory role that reflects preserved epithelial differentiation in HNSCC^40^. *TEAD3*, together with *TEAD1*, is essential for maintaining basal keratinocyte proliferation and suppressing terminal differentiation; inhibition of TEAD activity leads to rapid activation of differentiation programs in human epidermis^41^. *MPZL2* mediates cell-cell adhesion in epithelial tissues^42^.

We applied several other machine learning methods to build, train and test models on the same dataset. Random Forest regression achieved a reasonable LOOCV fit (MAE = 20.17, R = 0.69) using only 10 genes (*JPH2, CRIP1, HTR2C, ANOS1, ANXA8, MUC15, KLF4, C11orf71, CPVL, KRT15*) and exhibited strong performance on the clinical test set (MAE = 11.84, R = 0.73; **Fig. S3A, S3B**). Among the selected features were *KLF4* and *KRT15*, genes linked to differentiation and epithelial identity^43,44^. Using Elastic Net regression, the training fit was modest (MAE = 21.39, R = 0.33 with 25 genes; **Fig. S3C**) but performance improved on the test set (MAE = 9.33, R = 0.79) (**Fig. S3D**). Key predictors included *MKI67*, an established proliferation marker and *SCD5*, a stearoyl-CoA desaturase linked to epithelial differentiation^45,46^. XGBoost regression showed moderate performance (CV MAE = 18.21, CV R = 0.35; test MAE = 17.68, test R = 0.42; **Fig. S3E, S3F**) using a 38-gene model, which included *WNT2B*, a gene implicated in EGFR-WNT signaling cross-talk, particularly in modulating epithelial plasticity and resistance mechanisms^47^.

Together, these results demonstrate that erlotinib sensitivity in HPV-negative HNSCC can be robustly predicted from transcriptomics features. By integrating machine learning with patient-derived 3D microtumors, we developed and validated predictive models that generalize to clinical samples and reveal biologically meaningful gene signatures.

## DISCUSSION

HNSCC remains a biologically diverse and clinically challenging disease in which molecular stratification has not yet delivered therapeutic precision. Here, we developed an integrated approach that connects tumor transcriptomic state to functional dependencies, drug sensitivities, and clinically relevant predictors of targeted therapy response. Our analysis resolves HPV-negative HNSCC into mechanistically distinct tumor states that transcend traditional “basal” and “classical” labels. By harmonizing five independent RNA-seq datasets, we identify subgroups characterized by proliferative signaling (MYC, MET/FAK), epithelial differentiation and adhesion programs, EMT-like features with stromal activation, and mitochondrial metabolic wiring. These states reflect known biological axes in HNSCC yet provide greater resolution and functional interpretability than previous taxonomies. Within this spectrum, a proliferative axis is marked by MYC activity, MET/FAK signaling, inflammatory programs, and translational machinery. MYC is frequently upregulated in HNSCC and linked to aggressive growth, therapeutic resistance, and modulation of the tumor-immune interface ^48^, while MET and FAK promote proliferation, invasion, and treatment resistance, and functionally intersect with EGFR signaling to support aggressive disease phenotypes^49^. Co-activation of NF-κB, common in HNSCC, further reinforces survival and inflammatory signaling in this state ^50^. In contrast, a second axis reflects a well-differentiated epithelial program, enriched for keratinocyte differentiation, epidermal development, and cornified-envelope genes. Maintenance of epithelial adhesion and cell-cell cohesion, including E-cadherin expression, is associated with more differentiated histology and improved outcomes in HNSCC, providing disease-specific context for this state ^51^. These distinct transcriptional identities are not simply descriptive; they correspond to specific viability requirements and therapeutic vulnerabilities when overlaid with genome-scale CRISPR dependencies, underscoring that transcriptional state is tightly linked to dependency architecture and implying different therapeutic entry points within HPV-negative tumors.

Second, integrating CRISPR essentiality data with these transcriptional states demonstrates that subtype identity corresponds to distinct survival circuitry. While HPV-negative tumors share a core set of viability genes involved in RNA processing, ribosome function, DNA replication/repair, and cell-cycle surveillance ^52^, each subtype exhibits selective dependencies. The proliferative state relies on autophagy and mitotic machinery, consistent with prior evidence that autophagy supports HNSCC growth, and that AURKA-driven spindle control contributes to poor outcomes ^53^. The well-differentiated epithelial state depends on ERBB/PI3K signaling and cadherin-mediated adhesion, reflecting preserved epithelial programs. The mitochondrial-metabolic state shows reliance on oxidative phosphorylation and mitochondrial translation, in line with reports that OXPHOS sustains survival and therapy tolerance in HNSCC^54^. The EMT-enriched state exhibits dependencies in G2/M regulation, glycolysis, angiogenic signaling, integrin pathways, and Notch activity, consistent with invasive and stromal-interactive biology ^55,56^. Thus, transcriptional state maps to discrete functional liabilities, converting subtype distinctions into actionable biological dependencies.

A central feature of this study is translation of computational subtype biology into a clinically deployable tool. We developed and validated a transcriptomic predictor of EGFR inhibitor response using patient-derived 3D microtumors, a platform that retains native architecture and microenvironmental context. The ability of the model to generalize from cell lines to primary tumor tissue represents an important demonstration of feasibility for precision oncology in HNSCC. While EGFR expression alone has historically been insufficient to guide therapy in this disease^57^, our approach leverages broader transcriptional context to identify tumors with preserved epithelial differentiation programs and susceptibility to EGFR blockade. This provides a biologically grounded rationale for prospective biomarker development and for reevaluating EGFR-targeted strategies in molecularly selected patients.

Taken together, this work converts transcriptomic heterogeneity in HNSCC into an actionable map that links tumor states to what they depend on and how they respond to drugs and then turns that map into a predictor in patient tissue. Many solid tumors have recognized transcriptomic classes but lack actionable frameworks; our approach offers a template for converting tumor heterogeneity into treatment hypotheses that can be tested in ex vivo systems and, ultimately, in clinical trials. This framework is immediately extensible to other cancer types where descriptive molecular subgroups exist but are not yet clinically actionable, offering a systematic route to realizing the promise of precision oncology in biologically heterogeneous tumors.

## MATERIALS AND METHODS

### Study participant details

Human HNSCC tumor samples were obtained under IRB-approved protocols (#00001852 [STUDY00001852]). Tissues were collected with informed consent and de-identified prior to use **(Table S6)**.

### RNA extraction and library preparation

Total RNA was extracted from human tissue using the AllPrep DNA/RNA Mini Kit (Qiagen, 80204) per the manufacturer’s protocol. Fresh or snap-frozen samples were homogenized in RLT buffer with ceramic beads (Omni International, Cat. No. 19-646-3) using the Bead Ruptor 12 at 3-4 M/s for 5 seconds (1-2 cycles). Following a centrifugation, the resulting supernatant was processed through the kit’s spin columns to sequentially purify genomic DNA and total RNA. RNA was eluted in RNase-free water, and RNA integrity was assessed using the Agilent TapeStation. Samples were stored at -80°C. Library preparation, poly(A) enrichment, and paired-end (150 bp) sequencing were performed by Novogene on an Illumina HiSeq 2500.

### RNA sequencing data collection

RNA sequencing data for HNSCC was obtained from four different publicly available datasets. Raw reads were obtained from three GEO datasets - GSE74927, GSE205308, GSE178537. The raw RNAseq counts for TCGA data was downloaded from the GDC Portal via UCSC Xena Browser (https://xenabrowser.net/).

### RNAseq pre-processing

Raw SRA reads were first converted to FASTQ format using SRA-Toolkit. Quality control was performed on a total of 187 FASTQ files using FastQC (https://www.bioinformatics.babraham.ac.uk/projects/fastqc/) and MultiQC (https://github.com/MultiQC/MultiQC). Adapter trimming was applied to files as needed using Trimmomatic^58^. Using STAR^11^ and Samtools^59^, reads were aligned to the GRCh38.p13 genome and indexed. We filtered out any samples with fewer than 60% uniquely mapped reads. Raw counts were generated using featureCounts, a component of the Subread tool^60^. Batch correction was performed using ComBat-Seq^61^. Finally, we filtered the dataset to retain only protein-coding genes, using the GENCODE v40 annotation.

### Clustering

For our UMAP analysis, we transformed raw counts into variance-stabilizing transformed (VST) data using the ‘DESeq2’ package^62^. We then filtered the protein-coding genes to remove those with low variance (≤1) and low expression (≤1 in more than 20% of samples). The UMAP was constructed using ∼3,000 genes and 727 samples.

We then performed principal component analysis (PCA) using the ‘scikit-learn’ package, converting the count data into principal components (PCs). The top 20 PCs, accounting for 100% of the explained variance, were retained for further analysis. For UMAP construction, we used the ‘Manhattan’ distance metric, 16 nearest k-neighbors, a minimum distance of 0.11, and a spread of 5 (random seed 42).

To identify clusters, we applied four different clustering methods: HDBSCAN, Gaussian Mixture Models (GMM), Hierarchical Clustering, and k-means clustering. HDBSCAN was performed using the HDBSCAN tool with the Euclidean distance metric, a minimum of 4 samples, and a minimum cluster size of 20. The optimal number of clusters (k) was determined using four methods: Elbow Method, Silhouette Method, Davies-Bouldin Index (DBI), and Calinski-Harabasz Index (CHI). The optimal cluster number was then applied to all clustering methods. All clustering analyses, except for DBSCAN, were conducted using scikit-learn.

### Differential expression analysis

Differential expression analysis was performed using the ‘edgeR’ package^63^. Non-tumor samples were excluded from all group comparisons. Gene counts were first filtered using the ‘filterByExpr’ function before normalizing for library size and performing comparisons using the Exact Binomial Negative Test. Differentially expressed genes (DEGs) were defined as those with an absolute fold change > 1.5, FDR < 0.05, and minimum average CPM of 1.

### Enrichment analysis

Enrichment analysis was performed using GSEA-Preranked with ranks defined as (sign) fold change × (−log10(FDR)) and 1,000 permutations to compute Normalized Enrichment Scores. or subtype characterization, the top 50 up- and downregulated pathways per cluster (FDR < 25%) were retained.

ssGSEA 2.0 was applied to shortlisted gene sets using Transcripts Per Kilobase Million (TPM) values derived from GENCODE v40 gene lengths. CRISPR gene enrichment was analyzed with EnrichR in R using KEGG, Biocarta, Reactome, Hallmark, WikiPathways, and Gene Ontology (GO) databases^64^.

### Survival analysis

Kaplan-Meier survival analysis was performed to evaluate overall survival differences across HNSCC subtypes. Overall survival data was used for the analysis, with non-tumor samples excluded. To ensure consistency in comparisons, survival time was truncated at 60 months. Patients who died beyond 60 months were censored and assigned a status of “living”. Kaplan-Meier survival curves were generated using the “survival” R package, and statistical significance was assessed using the log-rank test.

### Cell deconvolution

For deconvolution, we used the xCell^65^ webtool and the default 64 immune and stromal subtype gene signatures.

### CRISPR gene effect scores

We collected the z-normalized gene effect scores for cell lines of our interest from DepMap (https://depmap.org/portal/).

### Drug response modeling

Gene expression (log₂(TPM+1)) from CCLE HNSC cell lines was filtered for genes with mean ≥ 0.5, variance ≥ 1, expression ≥ 1 in ≥ 5 samples, and variance-over-mean ≥ 0.1. These features were paired with experimentally derived erlotinib response values (0-100% viability; 100 = no effect). CCLE and clinical tumor samples (n = 33) were batch-corrected with ComBat-seq. Models were exclusively trained on CCLE data and evaluated on patient samples.

Hyperparameters for Random Forest and XGBoost were optimized with Optuna v3.0.3^66^. Recursive feature elimination (RFE) was used to identify the optimal number of predictors. Model performance was evaluated using mean absolute error (MAE) and the Pearson correlation coefficient (R).

*SVM:* SVR models were trained on standardized data using the RobustScaler and a linear kernel with C = 1.

*Elastic Net:* Same as SVR, with regularization strength (α) = 0.01 and L1 ratio = 0.8.

*XGBoost:* No scaling beyond log₂(TPM+1); parameters tuned: estimators [10-500], depth [2-10], learning rate [0.01-0.3], subsample [0.6-1.0], colsample_bytree [0.4-1.0]

*Random Forest:* Scaled using a QuantileTransformer prior to model fitting. Hyperparameters were optimized with MAE as the objective function. Parameters tuned: number of estimators [10, 200], maximum tree depth [2, 20], and maximum features considered at each split [’sqrt’, ’log2’, 0.1, 0.3, 0.5, 0.7]

### *3D* microtumors preparation and drug screening

Microtumors were prepared from HNSCC surgical resections as described previously ^67^. Briefly, tumor slices were prepared using a Leica VT1200S vibratome (Leica Biosystems, Wetzlar, Germany). To create microtumors, slices (400 μm thickness, 6 mm in diameter) were arranged on a plastic disc and sectioned using a McIlwain Tissue Chopper (Ted Pella, Redding, CA, USA). 3D microtumors were cultured in Williams’ Medium E containing 12 mM nicotinamide, 150 nM ascorbic acid, 2.25 mg/mL sodium bicarbonate, 20 mM HEPES, 50 mg/mL glucose, 1 mM sodium pyruvate, 2 mM L-glutamine, 1% (v/v) ITS, 20 ng/mL EGF, 40 IU/mL penicillin and 40 μg/mL streptomycin. 3D microtumors were treated with Realtime-Glo MT cell viability reagent (Promega, Madison, WI, USA) at 1: 1000 dilution in 96-well plates. Following a 24-h incubation, luminescence signals were measured using a Biotek Synergy H1 microplate reader (Agilent Technologies, Santa Clara, CA, USA). 3D microtumors were maintained and real-time viability was measured for 3 to 7 days post-treatment. The viability was determined by calculating the ratio of the signal at the end of the study to the signals from the same tissue before treatment, which was then normalized to the value of vehicle control 3D microtumors set to 100%.

### Data Visualization

All the visualization for the manuscript was done on R packages ggplot2, pheatmap and python’s matplotlib and Seaborn.

### Immunohistochemistry

HNSCC tumor tissues were fixed in 10% formalin and embedded into paraffin block to prepare tissue sections. Hematoxylin and Eosin (H&E), Masson’s Trichrome, Ki67, and CD45 staining were performed by the Fred Hutchinson Cancer Center Experimental Histopathology shared resource. For visualization, the images of entire tissue sections were obtained using the TissueFAXS system (TissueGnostics, Austria) with a 10x objective lens in brightfield mode.

### Materials availability

This study did not generate new unique reagents.

### Data and code availability

- The raw RNA-seq data has been provided in Supplementary data
- This paper does not report original code.
- Any additional information required to reanalyze the data reported in this work paper is available from the lead contact upon request.

## Acknowledgments

We thank Yuqi Kang for assistance with modeling. This research was supported by the Experimental Histopathology Shared Resources of the Fred Hutch/University of Washington Cancer/Seattle Children’s Cancer Consortium (P30 CA015704).

## Author Contributions

J.V. and T.S.G. conceived the study. J.V. performed all computational analysis and wrote the original draft. S.Z. and M.C. performed experiments. M.U., L.S., E.M., C.R. and B.B. provided clinical resources. S.B., B.C. and T.S.G. supervised the study and acquired funding.

---

**Fig. S1.**
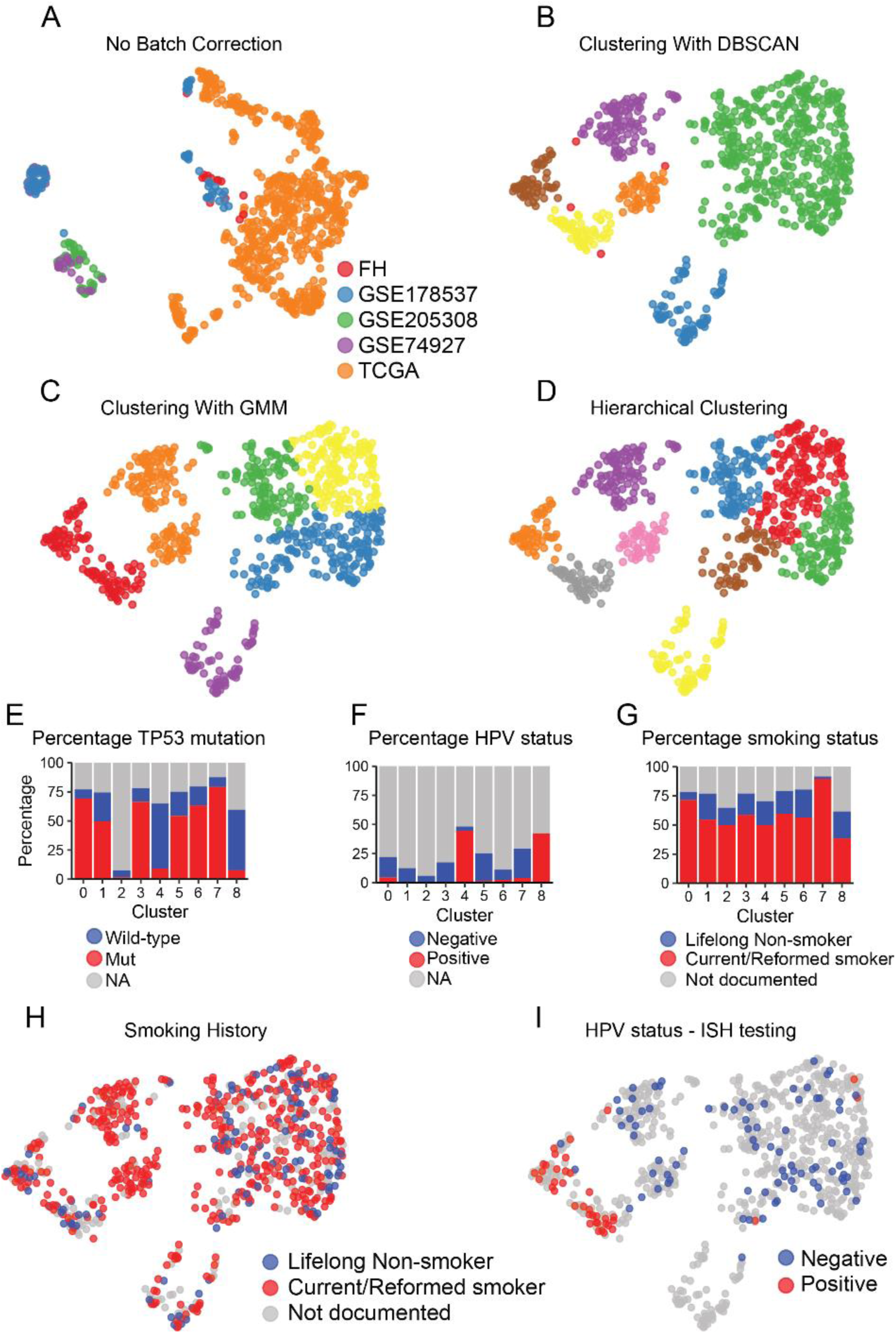
related to Fig. 1, 2. Comparative clustering and clinical annotations of HNSCC. **A** UMAP of integrated HNSCC samples without batch correction, with colors denoting dataset of origin (FHCC, GEO datasets, TCGA). **B** UMAP of HNSCC samples clustered using DBSCAN, with colors indicating distinct clusters. **C** UMAP of HNSCC samples clustered using Gaussian Mixture Modeling (GMM), showing separation into distinct groups. **D** UMAP of HNSCC samples clustered using hierarchical clustering, with each color representing a cluster. **E** Bar plots showing the percentage of TP53 mutation status (wild-type, mutant, NA) across clusters. **F** Bar plots showing the percentage of HPV status (positive, negative, NA) across clusters. **G** Bar plots showing the percentage of smoking status (lifelong non-smoker, current/reformed smoker, not documented) across clusters. **H** UMAP indicating smoking history, with samples colored by smoking status. **I** UMAP indicating HPV status from ISH testing, with samples colored by HPV-positive or -negative classification.

**Fig. S2.**
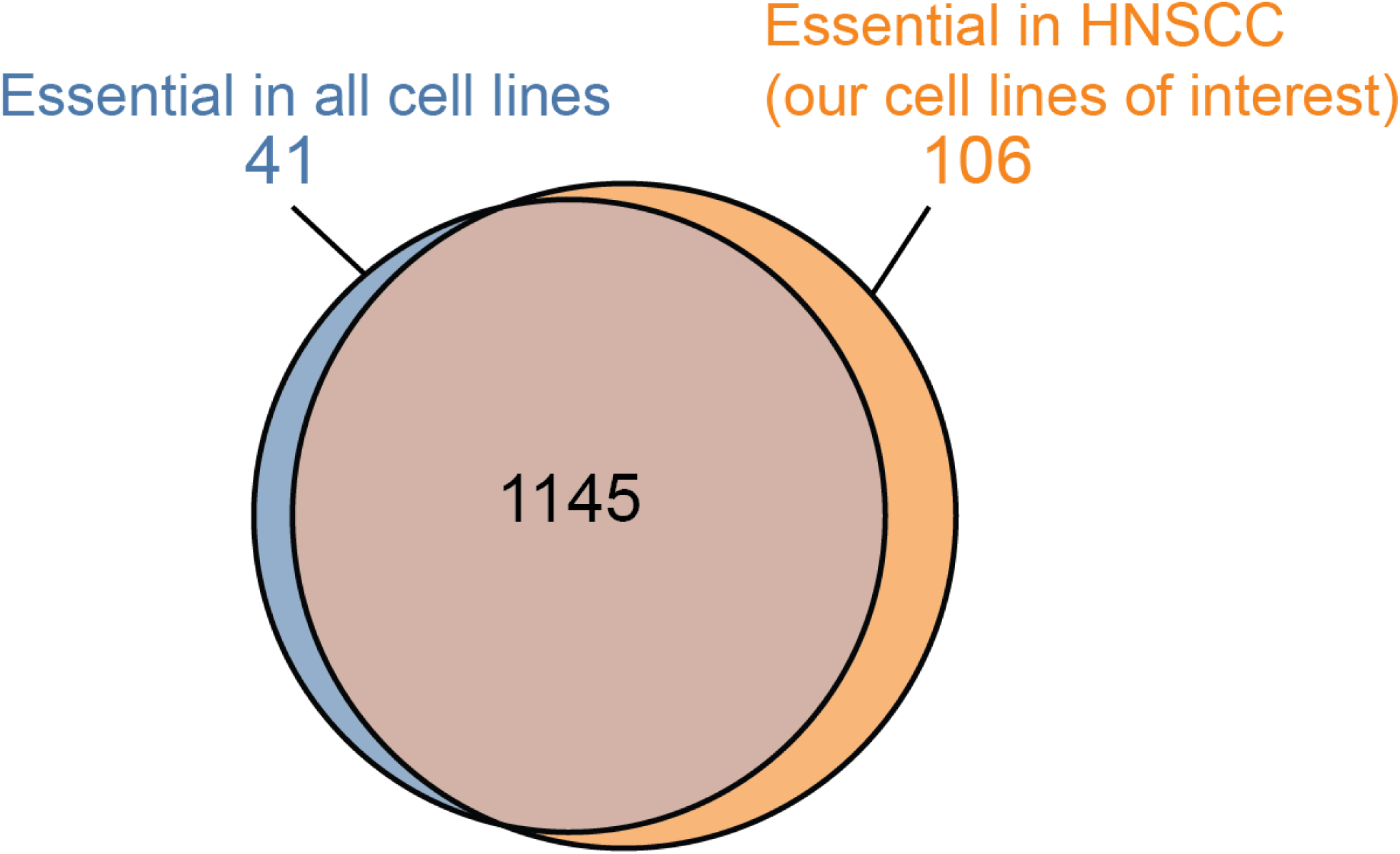
related to Fig. 4. Overlap of CRISPR-defined essential genes in HNSCC vs. other cancers. Venn diagram showing genes essential in HNSCC (n=18) and all cell lines (n=1178). Essentiality is defined as present in ≥80% of all cell lines, matching the essentiality threshold used in the HNSCC subtype-specific analysis.

**Fig. S3.**
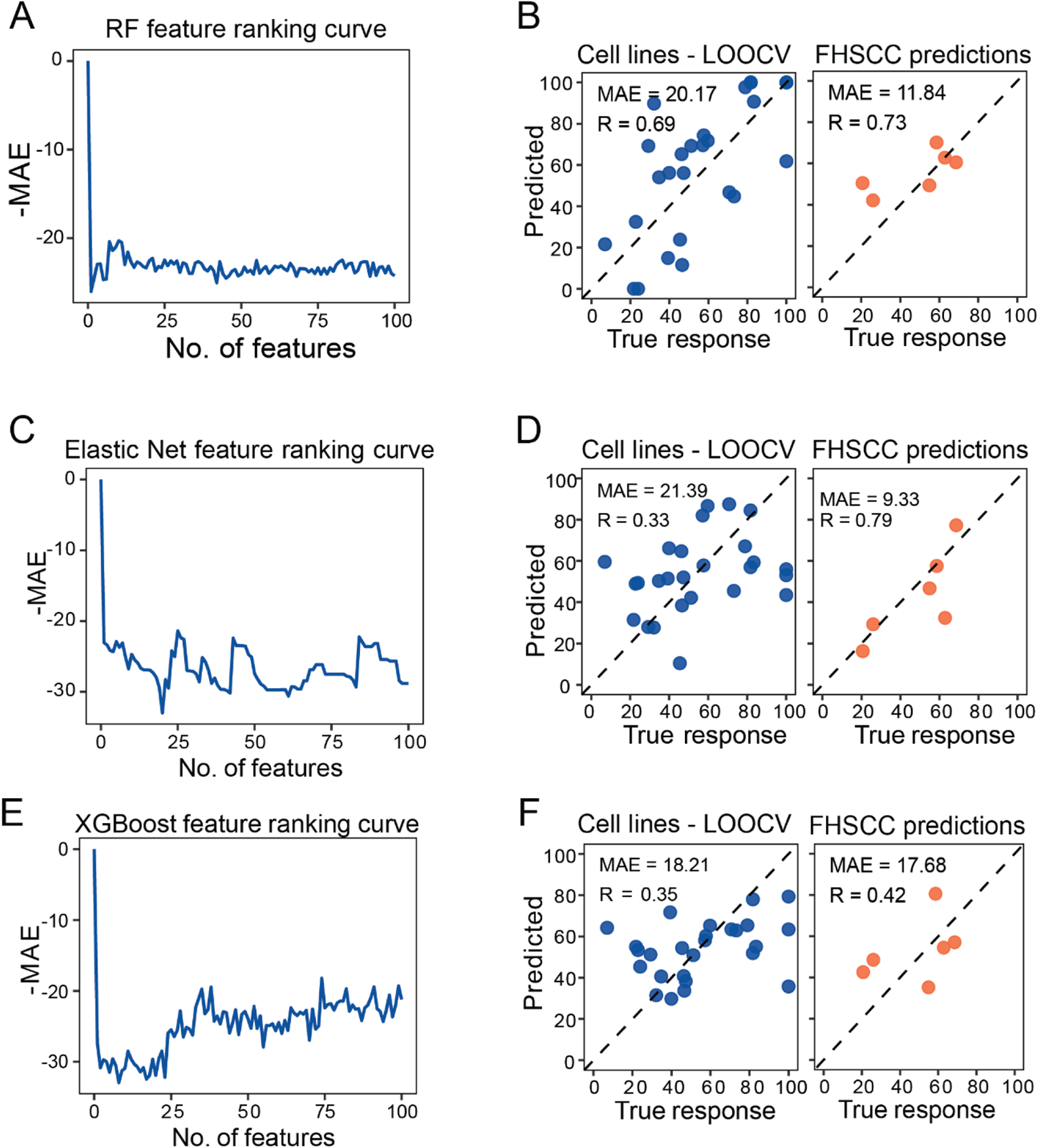
related toFig. 6. Comparative performance of machine learning models for predicting erlotinib sensitivity. **A** Feature ranking curve showing cross-validation MAE versus number of features for the Random Forest model. **B** Scatter plots of predicted versus true erlotinib responses for cell lines (left, LOOCV) and FHCC patient samples (right) using the Random Forest regression model. **C** Feature ranking curve showing cross-validation MAE versus number of features for the Elastic Net model. **D** Scatter plots of predicted versus true erlotinib responses for cell lines (left, LOOCV) and FHCC patient samples (right) using the Elastic Net regression model. **E** Feature ranking curve showing cross-validation MAE versus number of features for the XGBoost model. **F** Scatter plots of predicted versus true erlotinib responses for cell lines (left, LOOCV) and FHCC patient samples (right) using the XGBoost regression model.

## REFERENCES

1 Bray, F. et al. Global cancer statistics 2022: GLOBOCAN estimates of incidence and mortality worldwide for 36 cancers in 185 countries. Ca-Cancer J Clin 74, 229–263 (2024). 10.3322/caac.21834

2 Chung, C. H. et al. Molecular classification of head and neck squamous cell carcinomas using patterns of gene expression. Cancer cell 5, 489–500 (2004).

3 Lawrence, M. S. et al. Comprehensive genomic characterization of head and neck squamous cell carcinomas. Nature 517, 576–582 (2015). 10.1038/nature14129

4 Cheng, H. et al. Genomic and Transcriptomic Characterization Links Cell Lines with Aggressive Head and Neck Cancers. Cell Reports 25, 1332–1345.e1335 (2018). 10.1016/j.celrep.2018.10.007

5 Gu, Z. et al. Pharmacogenomic landscape of head and neck squamous cell carcinoma informs precision oncology therapy. Science Translational Medicine 14, eabo5987 (2022). doi:10.1126/scitranslmed.abo5987

6 Keck, M. K. et al. Integrative analysis of head and neck cancer identifies two biologically distinct HPV and three non-HPV subtypes. Clinical cancer research 21, 870–881 (2015).

7 Walter, V. et al. Molecular Subtypes in Head and Neck Cancer Exhibit Distinct Patterns of Chromosomal Gain and Loss of Canonical Cancer Genes. PLOS ONE 8, e56823 (2013). 10.1371/journal.pone.0056823

8 Vermorken, J. B. et al. Platinum-based chemotherapy plus cetuximab in head and neck cancer. New Engl J Med 359, 1116–1127 (2008). 10.1056/NEJMoa0802656

9 De Cecco, L. et al. Head and neck cancer subtypes with biological and clinical relevance: Meta-analysis of gene-expression data. Oncotarget 6, 9627 (2015).

10 Heath, A. P. et al. The NCI genomic data commons. Nature genetics 53, 257–262 (2021).

11 Dobin, A. et al. STAR: ultrafast universal RNA-seq aligner. Bioinformatics 29, 15–21 (2013).

12 Zhang, Y., Parmigiani, G. & Johnson, W. E. ComBat-seq: batch effect adjustment for RNA-seq count data. NAR genomics and bioinformatics 2, lqaa078 (2020).

13 Poeta, M. L. et al. TP53 Mutations and Survival in Squamous-Cell Carcinoma of the Head and Neck. New Engl J Med 357, 2552–2561 (2007). 10.1056/NEJMoa073770

14 Johnson, D. E. et al. Head and neck squamous cell carcinoma. Nature Reviews Disease Primers 6, 92 (2020). 10.1038/s41572-020-00224-3

15 Li, H. et al. Association of Human Papillomavirus Status at Head and Neck Carcinoma Subsites With Overall Survival. JAMA Otolaryngology–Head & Neck Surgery 144, 519–525 (2018). 10.1001/jamaoto.2018.0395

16 Lee, S. et al. SOX2 regulates self-renewal and tumorigenicity of stem-like cells of head and neck squamous cell carcinoma. British journal of cancer 111, 2122–2130 (2014).

17 Sriram, R. S. et al. Genetic alterations in CDKN2A interacting network and their putative association with head and neck squamous cell carcinoma. Human Gene 40, 201276 (2024).

18 Rampias, T., Sasaki, C. & Psyrri, A. Molecular mechanisms of HPV induced carcinogenesis in head and neck. Oral oncology 50, 356–363 (2014).

19 Subramanian, A. et al. Gene set enrichment analysis: a knowledge-based approach for interpreting genome-wide expression profiles. Proceedings of the National Academy of Sciences 102, 15545–15550 (2005).

20 Lawrence, M. S. et al. Comprehensive genomic characterization of head and neck squamous cell carcinomas. Nature 517, 576–582 (2015). 10.1038/nature14129

21 Aran, D., Hu, Z. & Butte, A. J. xCell: digitally portraying the tissue cellular heterogeneity landscape. Genome biology 18, 1–14 (2017).

22 Wang, Z. et al. Characterization of immune microenvironment in patients with HPV-positive and negative head and neck cancer. Scientific Data 10, 694 (2023). 10.1038/s41597-023-02611-3

23 Ghandi, M. et al. Next-generation characterization of the Cancer Cell Line Encyclopedia. Nature 569, 503–508 (2019). 10.1038/s41586-019-1186-3

24 Corsello, S. M. et al. Discovering the anticancer potential of non-oncology drugs by systematic viability profiling. Nature Cancer 1, 235–248 (2020). 10.1038/s43018-019-0018-6

25 Bonner, J. A. et al. Radiotherapy plus cetuximab for squamous-cell carcinoma of the head and neck. N Engl J Med 354, 567–578 (2006). 10.1056/NEJMoa053422

26 Traxler, P. et al. AEE788: A Dual Family Epidermal Growth Factor Receptor/ErbB2 and Vascular Endothelial Growth Factor Receptor Tyrosine Kinase Inhibitor with Antitumor and Antiangiogenic Activity. Cancer Res 64, 4931–4941 (2004). 10.1158/0008-5472.Can-03-3681

27 Carboni, J. M. et al. BMS-754807, a small molecule inhibitor of insulin-like growth factor-1R/IR. Molecular Cancer Therapeutics 8, 3341–3349 (2009). 10.1158/1535-7163.Mct-09-0499

28 Fletcher, G. C. et al. ENMD-2076 Is an Orally Active Kinase Inhibitor with Antiangiogenic and Antiproliferative Mechanisms of Action. Molecular Cancer Therapeutics 10, 126–137 (2011). 10.1158/1535-7163.Mct-10-0574

29 Spector, T., Lobe, D. C., Ellis, M. N., Blumenkopf, T. A. & Szczech, G. M. Inactivators of herpes simplex virus ribonucleotide reductase: hematological profiles and in vivo potentiation of the antiviral activity of acyclovir. Antimicrobial Agents and Chemotherapy 36, 934–937 (1992). 10.1128/aac.36.5.934

30 Rew, Y. & Sun, D. Discovery of a Small Molecule MDM2 Inhibitor (AMG 232) for Treating Cancer. Journal of Medicinal Chemistry 57, 6332–6341 (2014). 10.1021/jm500627s

31 Nakajima, T. et al. Y-27152, a long-acting K+ channel opener with less tachycardia: antihypertensive effects in hypertensive rats and dogs in conscious state. The Journal of Pharmacology and Experimental Therapeutics 261, 730–736 (1992). 10.1016/S0022-3565(25)11098-7

32 Li, C. J., Wang, C. & Pardee, A. B. Induction of Apoptosis by β-Lapachone in Human Prostate Cancer Cells1. Cancer Res 55, 3712–3715 (1995).

33 Cruz-Gregorio, A., Martínez-Ramírez, I., Pedraza-Chaverri, J. & Lizano, M. Reprogramming of Energy Metabolism in Response to Radiotherapy in Head and Neck Squamous Cell Carcinoma. Cancers 11, 182 (2019).

34 Aronne, L. J. et al. Efficacy and Safety of CP-945,598, a Selective Cannabinoid CB1 Receptor Antagonist, on Weight Loss and Maintenance. Obesity 19, 1404–1414 (2011). 10.1038/oby.2010.352

35 Kalyankrishna, S. & Grandis, J. R. Epidermal growth factor receptor biology in head and neck cancer. J Clin Oncol 24, 2666–2672 (2006). 10.1200/jco.2005.04.8306

36 Wee, P. & Wang, Z. Epidermal Growth Factor Receptor Cell Proliferation Signaling Pathways. Cancers (Basel) 9 (2017). 10.3390/cancers9050052

37 Kang, Y., Vijay, S. & Gujral, T. S. Deep neural network modeling identifies biomarkers of response to immune-checkpoint therapy. iScience 25, 104228 (2022). 10.1016/j.isci.2022.104228

38 Chan, M. et al. DNAJ-PKAc fusion heightens PLK1 inhibitor sensitivity in fibrolamellar carcinoma. Gut (2025).

39 Nishida-Aoki, N., Bondesson, A. J. & Gujral, T. S. Measuring Real-time Drug Response in Organotypic Tumor Tissue Slices. J Vis Exp (2020). 10.3791/61036

40 Novotný, J. et al. Analysis of HPV-Positive and HPV-Negative Head and Neck Squamous Cell Carcinomas and Paired Normal Mucosae Reveals Cyclin D1 Deregulation and Compensatory Effect of Cyclin D2. Cancers 12, 792 (2020).

41 Yuan, Y. et al. YAP1/TAZ-TEAD transcriptional networks maintain skin homeostasis by regulating cell proliferation and limiting KLF4 activity. Nature Communications 11, 1472 (2020). 10.1038/s41467-020-15301-0

42 Wesdorp, M. et al. MPZL2, Encoding the Epithelial Junctional Protein Myelin Protein Zero-like 2, Is Essential for Hearing in Man and Mouse. Am J Hum Genet 103, 74–88 (2018). 10.1016/j.ajhg.2018.05.011

43 Giroux, V. et al. Long-lived keratin 15+ esophageal progenitor cells contribute to homeostasis and regeneration. J Clin Invest 127, 2378–2391 (2017). 10.1172/jci88941

44 Han, J. et al. Identification of potential therapeutic targets in human head & neck squamous cell carcinoma. Head Neck Oncol 1, 27 (2009). 10.1186/1758-3284-1-27

45 Puglisi, R. et al. SCD5 restored expression favors differentiation and epithelial-mesenchymal reversion in advanced melanoma. Oncotarget 9, 7567–7581 (2018). 10.18632/oncotarget.24085

46 Shirima, C. A. et al. Epithelial-derived head and neck squamous tumourigenesis (Review). Oncol Rep 52 (2024). 10.3892/or.2024.8800

47 Hu, T. & Li, C. Convergence between Wnt-β-catenin and EGFR signaling in cancer. Mol Cancer 9, 236 (2010). 10.1186/1476-4598-9-236

48 Field, J. et al. Elevated expression of the c-myc oncoprotein correlates with poor prognosis in head and neck squamous cell carcinoma. Oncogene 4, 1463–1468 (1989).

49 Eke, I. & Cordes, N. Dual targeting of EGFR and focal adhesion kinase in 3D grown HNSCC cell cultures. Radiother Oncol 99, 279–286 (2011). 10.1016/j.radonc.2011.06.006

50 Ondrey, F. G. et al. Constitutive activation of transcription factors NF-κB, AP-1, and NF-IL6 in human head and neck squamous cell carcinoma cell lines that express pro-inflammatory and pro-angiogenic cytokines. Molecular Carcinogenesis 26, 119–129 (1999). 10.1002/(SICI)1098-2744(199910)26:2<119::AID-MC6>3.0.CO;2-N

51 Schipper, J. H. et al. E-cadherin expression in squamous cell carcinomas of head and neck: inverse correlation with tumor dedifferentiation and lymph node metastasis. Cancer Res 51, 6328–6337 (1991).

52 Huang, C. et al. Proteogenomic insights into the biology and treatment of HPV-negative head and neck squamous cell carcinoma. Cancer Cell 39, 361–379.e316 (2021). 10.1016/j.ccell.2020.12.007

53 Rikiishi, H. Autophagic action of new targeting agents in head and neck oncology. Cancer Biol Ther 13, 978–991 (2012). 10.4161/cbt.21079

54 Koc, E. C. et al. Impaired mitochondrial protein synthesis in head and neck squamous cell carcinoma. Mitochondrion 24, 113–121 (2015). 10.1016/j.mito.2015.07.123

55 Inamura, N. et al. Notch1 regulates invasion and metastasis of head and neck squamous cell carcinoma by inducing EMT through c-Myc. Auris Nasus Larynx 44, 447–457 (2017). 10.1016/j.anl.2016.08.003

56 Jumaniyazova, E., Lokhonina, A., Dzhalilova, D., Kosyreva, A. & Fatkhudinov, T. Role of Microenvironmental Components in Head and Neck Squamous Cell Carcinoma. J Pers Med 13 (2023). 10.3390/jpm13111616

57 Cooper, J. B. & Cohen, E. E. W. Mechanisms of resistance to EGFR inhibitors in head and neck cancer. Head & Neck 31, 1086–1094 (2009). 10.1002/hed.21109

58 Bolger, A. M., Lohse, M. & Usadel, B. Trimmomatic: a flexible trimmer for Illumina sequence data. Bioinformatics 30, 2114–2120 (2014). 10.1093/bioinformatics/btu170

59 Li, H. et al. The Sequence Alignment/Map format and SAMtools. Bioinformatics 25, 2078–2079 (2009). 10.1093/bioinformatics/btp352

60 Liao, Y., Smyth, G. K. & Shi, W. featureCounts: an efficient general purpose program for assigning sequence reads to genomic features. Bioinformatics 30, 923–930 (2014). 10.1093/bioinformatics/btt656

61 Zhang, Y., Parmigiani, G. & Johnson, W. E. ComBat-seq: batch effect adjustment for RNA-seq count data. NAR Genom Bioinform 2, lqaa078 (2020). 10.1093/nargab/lqaa078

62 Love, M. I., Huber, W. & Anders, S. Moderated estimation of fold change and dispersion for RNA-seq data with DESeq2. Genome Biol 15, 550 (2014). 10.1186/s13059-014-0550-8

63 Robinson, M. D., McCarthy, D. J. & Smyth, G. K. edgeR: a Bioconductor package for differential expression analysis of digital gene expression data. Bioinformatics 26, 139–140 (2010). 10.1093/bioinformatics/btp616

64 Kuleshov, M. V. et al. Enrichr: a comprehensive gene set enrichment analysis web server 2016 update. Nucleic Acids Res 44, W90–97 (2016). 10.1093/nar/gkw377

65 Aran, D., Hu, Z. & Butte, A. J. xCell: digitally portraying the tissue cellular heterogeneity landscape. Genome Biol 18, 220 (2017). 10.1186/s13059-017-1349-1

66 Akiba, T., Sano, S., Yanase, T., Ohta, T. & Koyama, M. in *Proceedings of the 25th ACM SIGKDD International Conference on Knowledge Discovery & Data Mining* 2623–2631 (Association for Computing Machinery, Anchorage, AK, USA, 2019).

67 Nishida-Aoki N, Z. S., Chan M, Kang Y, Fujita M, Jiang X, McCabe M, Vaz JM, Davidson NE, Ghajar CM, Hansen K, Welm AL, Pillarisetty VG, Gujral TS. Drug screening in 3D microtumors reveals DDR1/2-MAPK12-GLI1 as a vulnerability in cancer-associated fibroblasts. Cell Reports Medicine In press (2025).

